# Genetic risk for neurodegenerative conditions is linked to disease-specific microglial pathways

**DOI:** 10.1101/2024.08.29.610255

**Authors:** Aydan Askarova, Reuben M. Yaa, Sarah J. Marzi, Alexi Nott

## Abstract

Genome-wide association studies have identified thousands of common variants associated with an increased risk of neurodegenerative disorders. However, the noncoding localization of these variants has made the assignment of target genes for brain cell types challenging. Genomic approaches that infer chromosomal 3D architecture can link noncoding risk variants and distal gene regulatory elements such as enhancers to gene promoters. By using enhancer-to-promoter interactome maps for microglia, neurons, and oligodendrocytes, we identified cell-type-specific enrichment of genetic heritability for brain disorders through stratified linkage disequilibrium score regression. Our analysis suggests that genetic heritability for multiple neurodegenerative disorders is enriched at microglial chromatin contact sites. Through Hi-C coupled multimarker analysis of genomic annotation (H-MAGMA) we identified disease risk genes for Alzheimer’s disease, Parkinson’s disease, multiple sclerosis and amyotrophic lateral sclerosis. We found that disease-risk genes were overrepresented in microglia compared to other brain cell types across neurodegenerative conditions. Notably, the microglial risk genes and pathways identified were largely specific to each disease. Our findings reinforce microglia as an important, genetically informed cell type for therapeutic interventions in neurodegenerative conditions and highlight potentially targetable disease-relevant pathways.

## Introduction

Genetics plays a significant role in the etiology of neurodegenerative disorders including Alzheimer’s disease (AD), Parkinson’s disease (PD), multiple sclerosis (MS) and amyotrophic lateral sclerosis (ALS) (2, 3, 4, 5, 6, 7, 8, 9, 10, 11). Familial forms have been identified for AD, PD and ALS that exhibit Mendelian patterns of inheritance and are associated with rare variants with strong effect sizes (12, 13, 14, 15, 16, 17, 18, 19, 20, 21, 22, 23, 24, 25, 26, 27). While the genetics underlying familial cases have been informative in our understanding of disease etiology, most individuals presenting with neurogenerative disorders, including MS, have sporadic forms of the disease. Genome-wide association studies (GWAS) for sporadic neurodegenerative disorders have identified thousands of common variants associated with an increased risk of disease and highlight the heterogeneity of these disorders (28, 29, 30, 31, 32, 33, 34). GWAS risk variants generally have a relatively high prevalence in the population and exhibit smaller effect sizes, with their risk contribution believed to arise from the combined effects of multiple variants (35).

Most GWAS risk variants reside within non-coding regions of the genome and are often located distally from the nearest known genes (36). GWAS risk variants are enriched at chromatin accessibility regions that likely function as gene regulatory elements such as enhancers and promoters (37, 38). Enhancers are distal genomic regions associated with chromatin accessibility and are characterized by the presence of specific histone modifications, including acetylation of histone H3 lysine 27 (H3K27ac) (39). Enhancers are highly cell type-specific (40) and can be incorporated into heritability analysis to prioritize cell types associated with the genetic risk of complex traits. GWAS variants for neurological disorders and psychiatric traits have been associated with cell type-specific heritability enrichment. For example, AD risk variants were found to be enriched in microglia and macrophage enhancers, while schizophrenia risk was enriched in neuronal gene regulatory regions (1, 41, 42, 43, 44, 45, 46, 47).

Enhancers have been informative for the allocation of cell types associated with genetic risk. However, the distal localization of GWAS risk variants has made the identification of target genes impacted by these variants a major challenge. The mammalian genome has a non-random three-dimensional organization that connects distal chromosomal regions through the formation of chromatin loops (48). Functional chromatin interactions include the association of gene promoters with *cis*-regulatory regions, such as enhancers (48). The recruitment of transcription factors and structural proteins to enhancers and their interaction with promoters facilitates the formation of the pre-initiation complex and gene transcription (49, 50). Genetic variants localized to gene regulatory regions were thought to disrupt enhancer function or enhancer-to-promoter interactions, ultimately impacting gene expression and cell behavior (46, 51). Similar to enhancers, chromatin interactions are cell-type-specific (1). Hence, localization of non-coding GWAS variants to chromatin contact sites could predict cell type-specific genes and pathways that are susceptible to genetic variation in neurodegenerative disorders.

Enhancer-to-promoter interactomes are available for three of the major brain cell types, however, the assignment of GWAS risk variants to genes has been hindered by a lack of computational tools. A recently developed tool, Hi-C coupled multimarker analysis of genomic annotation (H-MAGMA), identifies putative disease risk genes by accounting for GWAS variants within distal non-coding regions (52). H-MAGMA predicts gene-level associations with diseases by combining GWAS summary statistics with enhancer-to-promoter interactomes (52, 53). Here we used H-MAGMA coupled with chromatin data to map out disease genes for neurodegenerative diseases. By integrating epigenetic annotations with chromatin interaction data, we identified putative cell types and genes that contribute to the genetic susceptibility of these disorders. We found that risk genes are enriched in microglia across multiple neurodegenerative diseases (AD, PD, ALS, and MS). However, the pathways impacted by microglial GWAS-risk genes are mostly unique for each disorder, indicating that immune processes exhibit disease-specific patterns.

## Results

### Microglial chromatin interactions are enriched for genetic risk variants associated with neurodegenerative disorders

To determine whether disease risk variants for neurodegenerative disorders are associated with genes linked to distal gene regulatory regions we used proximity ligation-assisted chromatin immunoprecipitation-seq (PLAC-seq) data generated from human cortical neurons, microglia and oligodendrocytes (1). PLAC-seq chromatin interactions were anchored to active gene promoters by immunoprecipitation of histone H3 lysine 4 trimethylation (H3K4me3), which is a histone modification enriched at active gene promoters (54, 55). PLAC-seq contact sites were defined as two 5 kb regions (or bins) separated by 10 kb or more (1). By integrating PLAC-seq-defined chromatin loops with ATAC-seq, H3K27ac chromatin immunoprecipitation (ChIP)-seq and H3K4me3 ChIP-seq from the same cell types (1), we classified chromatin interactions as either: i) promoter-to-enhancer; ii) promoter-to-promoter; iii) promoter-to-ATAC; iv) promoter-to-promoter/enhancer; v) promoter-to-other; vi) H3K4me3-to-H3K4me3; vii) H3K4me3-to-other; and viii) other interactions. Active promoters were defined by co-occurrence of H3K4me3 and H3K27ac within 2 kb of a transcription start site (TSS). Enhancers were defined as H3K27ac peaks that did not overlap with H3K4me3. PLAC-seq contact sites that overlapped both promoter and enhancer regions were termed promoter/enhancer. Genomic regions with H3K4me3 peaks further than 2 kb from a TSS were not considered promoters and were classified as ‘H3K4me3’ regions. Distal regions that were linked to promoters and had an ATAC peak but no H3K27ac peak were defined as ‘ATAC’. Lastly, chromatin loops that linked to regions with no detectable H3K4me3, H3K27ac or ATAC signal were designated ‘other’. Promoter-to-enhancer loops were the most common classification of chromatin interactions for each cell type, representing 29.4%, 39.1% and 38.2% of interactions in microglia, neurons, and oligodendrocytes, respectively (**Fig. 1a**). The next most abundant classifications were chromatin interactions that occurred at promoters-to-other or H3K4me3-to-other (**Fig. 1a**). Promoters are known to interact with more than one enhancer. For microglia, neurons and oligodendrocytes, most promoters interacted with more than one enhancer and for enhancer-to-promoter interactions, most enhancers interacted with a single promoter (**Fig. 1b**). The average distance of these promoter-to-enhancer interactions was 175 kb for microglia, 200 kb for neurons and 150 kb for oligodendrocytes (**Fig. 1c**). Overall, H3K4me3-anchored PLAC-seq chromatin loops in microglia, neurons and oligodendrocytes predominantly identified promoters that were linked to multiple distal enhancers.

**Figure 1.**
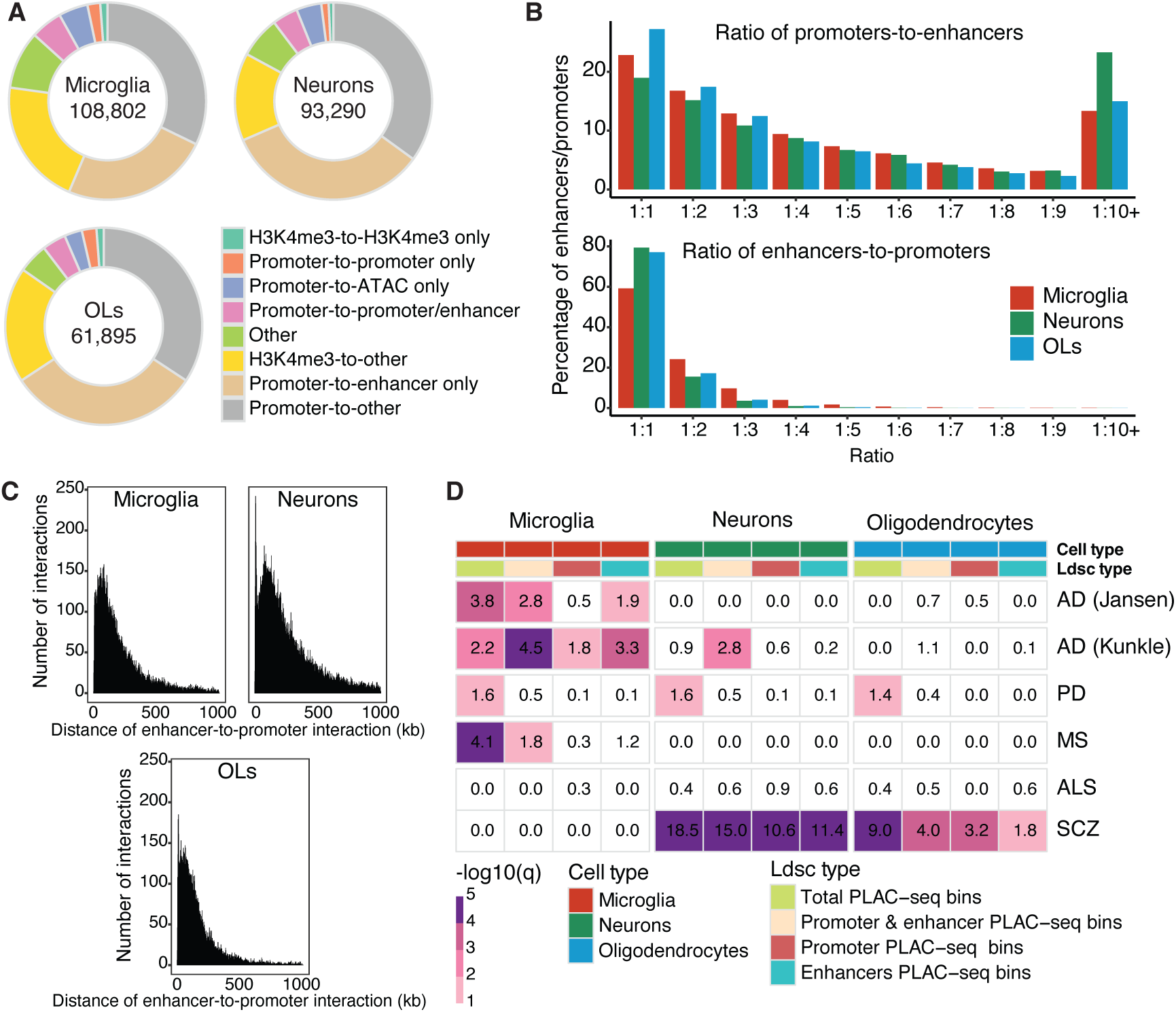
Microglia enhancer-to-promoter interactions were enriched for disease-risk variants across multiple neurodegenerative conditions. **A)** Doughnut plots of classifications of PLAC-seq interactions identified in human microglia, neurons and oligodendrocytes (1) with the total number of interactions shown in the center. ‘Promoters’, PLAC-seq bins that overlap a H3K4me3 and H3K27ac peak within 2,000 bp of a transcriptional start site (TSS). ‘Enhancers’, PLAC-seq bins that overlap H3K27ac peaks distal to the TSS. ‘H3K4me3’, PLAC-seq bins that overlap H3K4me3 peaks distal to TSS. ‘ATAC’, PLAC-seq bins that overlap chromatin accessible regions devoid of H3K4me3 and H3K27ac. **B)** Percent distribution of the number of enhancers interacting with individual promoters (top plot) and the number of promoters interacting with individual enhancers (bottom plot). **C)** Distribution plot of the proportion of distances between midpoints of promoters and midpoints of enhancers that interact based on chromatin interaction PLAC-seq data. **D)** Heatmap of partitioned heritability using sLDSC regression analysis of: (i) total PLAC-seq bins, (ii) promoter & enhancer PLAC-seq bins, (iii) promoter PLAC-seq bins and (iv) enhancer PLAC-seq bins for microglia, neurons and oligodendrocytes in AD (28, 29), PD (30) (excluding 23andMe), MS (34), ALS (32), and schizophrenia (33). Shown are LDSC enrichment p-values with Benjamini–Hochberg FDR correction for the number of diseases and cell types (-log10(q)). Disease enrichment was considered insignificant if the coefficient z-score was negative and assigned a 0.0 -log10(p) score. OLs, oligodendrocytes. SCZ, schizophrenia.

To examine whether disease heritability was enriched at brain cell type chromatin interactions, we used stratified linkage disequilibrium score (sLDSC) regression analysis. Cell type disease enrichment by sLDSC regression was assessed using chromatin interactions defined as (i) all PLAC-seq bins irrespective of functional genomic annotations (total PLAC-seq bins), (ii) PLAC-seq bins subset to both active gene promoters (H3K4me3 + H3K27ac) and distal enhancers (H3K27ac only) (promoter & enhancer PLAC-seq bins), (iii) PLAC-seq bins subset to active gene promoters (promoter PLAC-seq bins) and (iv) PLAC-seq bins subset to distal enhancers (enhancer PLAC-seq bins). Cell type disease enrichment was assessed using summary statistics from two complementary AD GWAS; one study was based exclusively on clinical diagnosis (28), while the second included by-proxy cases (29). GWAS summary statistics were analyzed for three additional neurodegenerative conditions, PD, MS and ALS (30, 32, 34) and for a neurodevelopmental condition, schizophrenia (33).

Microglia PLAC-seq bins showed enrichment for AD risk variants over other cell types, with a greater enrichment of AD risk variants found at enhancer PLAC-seq bins compared to promoter PLAC-seq bins (**Fig. 1d, Supplemental Fig. 1**). These findings corroborate observations that AD GWAS variants are enriched at microglia enhancers compared to microglia promoters defined using histone modifications (1, 56). The microglia enhancer PLAC-seq bins are likely physically linked to gene promoters and therefore functionally relevant. An enrichment of disease risk variants at microglia PLAC-seq bins was also observed for PD and MS, although there was no clear preference for either promoter or enhancer interacting regions for these disorders (**Fig. 1d, Supplemental Fig. 1**). No enrichment for ALS risk variants was identified at PLAC-seq bins for microglia, neurons or oligodendrocytes (**Fig. 1d, Supplemental Fig. 1**). In contrast, for schizophrenia, heritability showed a strong enrichment of disease risk at PLAC-seq bins identified in neurons and oligodendrocytes (**Fig. 1d, Supplemental Fig. 1**). The heritability enrichment for schizophrenia was stronger in neurons than oligodendrocytes (promoter & enhancer PLAC-seq bins; sLDSC; neurons -log10(q)=15; oligodendrocytes, -log10(q)=4.0) (**Fig. 1d**). This supports previous findings showing that schizophrenia GWAS variants were enriched at neuronal promoters and enhancers using annotations defined by histone modifications (1, 44). Overall, chromatin-interacting regions in microglia show a broad enrichment for disease heritability across multiple neurodegenerative disorders.

### Microglial chromatin interactions identify disease risk genes across multiple neurodegenerative conditions

Promoter-to-enhancer interactions link distal gene regulatory regions, such as enhancers, to active gene promoters and can be used to infer disease-risk genes for noncoding GWAS risk variants. H-MAGMA was used to identify disease-risk genes in microglia, neurons and oligodendrocytes for AD, PD, MS, ALS and schizophrenia by incorporating PLAC-seq interactomes for the corresponding cell types. In all the neurodegenerative GWAS that we assessed, the highest number of risk genes were identified in microglia compared to neurons and oligodendrocytes (**Fig. 2a, Supplementary Table 1**). In contrast, for schizophrenia the highest number of risk genes were identified in neurons (**Fig. 2a, Supplementary Table 1**). The identified number of PLAC-seq chromatin interactions were higher in microglia compared to other cell types (microglia, 108802; neurons, 93290; oligodendrocytes, 61895; **Fig. 1a**), which may partially explain the increased number of microglia disease risk genes identified across neurodegenerative conditions. To account for the differing number of chromatin interactions identified between the three cell types, the PLAC-seq data was randomly downsampled to 60,000 chromatin interactions per cell type. This was followed by H-MAGMA analysis, which was repeated for 10 iterations (**Fig. 2b**). H-MAGMA analysis using the 60,000 downsampled PLAC-seq chromatin interactions maintained a similar distribution of disease-risk genes across the three cell types (**Fig. 2b**). Importantly, when the number of chromatin interactions was the same for each cell type, the number of disease-risk genes identified remained highest in microglia for AD, PD, MS and ALS (**Fig. 2b**). The overrepresentation of disease risk genes identified in microglia for neurodegenerative disorders compared to neurons for schizophrenia is consistent with the cell type distribution of disease heritability identified using sLDSC regression analysis (**Fig. 1d**).

**Figure 2.**
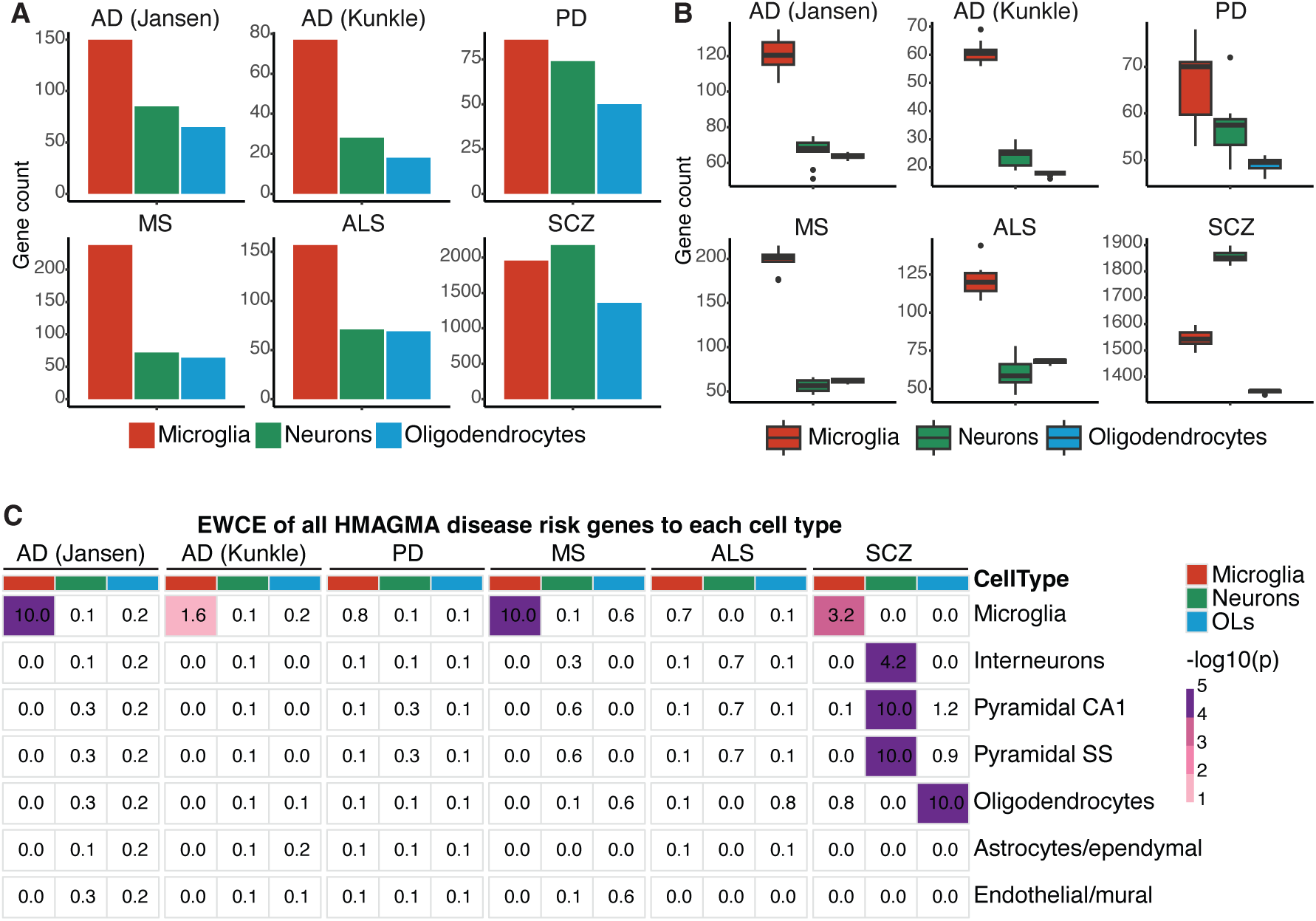
Microglial disease risk genes were identified for distal GWAS variants using chromatin loops. **A)** The number of disease risk genes identified in microglia, neurons and oligodendrocytes using H-MAGMA and GWAS for AD, PD (excluding 23andMe), MS, ALS, and schizophrenia. Gene-to-SNP associations were assigned for SNPs that were located within the promoter or exon of a gene, or within enhancers that were linked to genes through PLAC-seq interactions. **B)** To account for differences in chromatin interactions between cell types, the number of enhancer-to-promoter interactions was randomly sampled down to 60,000 loops 10 times. The number of disease risk genes were identified using the sampled down loops for microglia, neurons and oligodendrocytes with H-MAGMA for AD, PD (excluding 23andMe), MS, ALS, and schizophrenia. Dunn’s test (non-parametric) between cell types within each group: AD (Jansen 2019): microglia-neurons (**), microglia-oligo (****), neurons-oligo (ns); AD (Kunkle 2019): microglia-neurons (*), microglia-oligo (****), neurons-oligo (*); PD: microglia-neurons (ns), microglia-oligo (*****), neurons-oligo (**); MS: microglia-neurons (****), microglia-oligo (**), neurons-oligo (ns); ALS: microglia-neurons (****), microglia-oligo (**), neurons-oligo (ns); schizophrenia: microglia-neurons (*), microglia-oligo (*), neurons-oligo (****). **C)** EWCE analysis identified cell type enrichment of H-MAGMA disease risk genes from Fig. 2A using mouse cortex and hypothalamus single-cell RNA-seq (58). Shown are EWCE p-values. SCZ, schizophrenia.OLs, oligodendrocytes. *p <0.05, **p<0.01, ***p<1e-4, ****p<1e-6.

GWAS risk variants may be differentially enriched at chromatin interaction contact sites at enhancers (PLAC-seq bins at intergenic and intronic regions) compared to promoters (PLAC-seq bins at promoters and exonic regions). To determine GWAS risk enrichment across these gene regulatory classifications, H-MAGMA was repeated using PLAC-seq bins subset to enhancers or promoters. For AD, PD and MS, the maximum number of disease-risk genes were identified using microglia enhancer contact sites, followed by microglia promoter contact sites (**Supplemental Fig. 2**). In ALS, more disease-risk genes were identified using microglia promoter contact sites compared to enhancer contact sites (**Supplemental Fig. 2**). This suggests that promoters may play a more crucial role in the genetic risk associated with ALS, in contrast to the significance of enhancers for GWAS risk in other neurodegenerative conditions. Lastly, for schizophrenia, the highest number of disease-risk genes were identified at neuronal promoter contact sites compared to enhancers (**Supplemental Fig. 2**).

Disease-risk variants often colocalise with gene regulatory regions that are highly cell-type specific, thereby conferring cell-type-associated genetic susceptibility (37, 57). However, the downstream genes associated with these regulatory regions may be expressed exclusively in the disease-associated cell type or across multiple cell types. Expression Weighted Celltype Enrichment (EWCE) analysis was used to determine the cell type expression of the GWAS risk genes identified by H-MAGMA by incorporating single-cell gene expression data from the mouse cortex and hippocampus (58). EWCE analysis revealed that the expression of microglia GWAS-risk genes for AD, MS and schizophrenia was enriched in microglia compared to other brain cell types (**Fig. 2c**). In contrast, GWAS-risk genes identified in neurons and oligodendrocytes across the neurodegenerative conditions generally did not exhibit a cell type enrichment in gene expression, indicating a broader expression across multiple cell types (**Fig. 2c**). However, disease risk genes identified by H-MAGMA genes across all three cell types for schizophrenia were characterized by matching cell type-specific gene expression (**Fig. 2c**). Of note, neurons and oligodendrocytes originate from neural progenitor cells localized in the brain (59, 60), while microglia are derived from a distinct progenitor pool in the embryonic yolk sac outside of the brain (61). This may account for the cell type specificity in gene expression of microglia-associated risk genes across neurological conditions compared to risk genes identified in neurons and oligodendrocytes.

### Microglial genetic-susceptibility genes are associated with disease-specific pathways

Genetic heritability estimates using sLDSC and the identification of putative GWAS risk genes using H-MAGMA highlight the importance of microglia in the genetic susceptibility of neurodegenerative conditions. This may suggest shared dysregulated microglial processes across these disorders. However, an intersection of GWAS-risk genes identified using H-MAGMA for microglia showed a minimal overlap between the different diseases (filtered on H-MAGMA p-value; AD, PD, MS, ALS p<5e-8 and schizophrenia p<5e-12) (**Fig. 3a,b**). Similarly, there was a minimal overlap across diseases for GWAS risk genes identified for neurons and oligodendrocytes (**Fig. 3a,b**). While most risk genes were unique to each disorder, some genes were shared across two or more conditions. For example, the major histocompatibility complex (MHC) was identified as a disease-risk locus in MS and schizophrenia (**Fig. 3b**). Disease-risk genes that overlapped across PD, ALS and schizophrenia were *KANSL1-AS1* (microglia and oligodendrocytes) and *KANSL1*, *ARHGAP27*, and *PLEKHM1* (microglia). Interestingly, *KANSL1* and *ARHGAP27* were identified as comorbid genes for PD and ALS (62). The microglial GWAS-risk genes *BAG6*, *NEU1*, *PRRC2A*, *PSMB8*, *PSMB8-AS1* and *PSMB9* were associated with MS, ALS and schizophrenia. *PSMB8-AS1* was also identified as a microglial risk gene for AD. These findings indicate that microglia are an important cell type associated with genetic susceptibility across multiple neurodegenerative disorders. However, the microglial genes that are impacted by genetic risk are mostly disease-specific.

**Figure 3.**
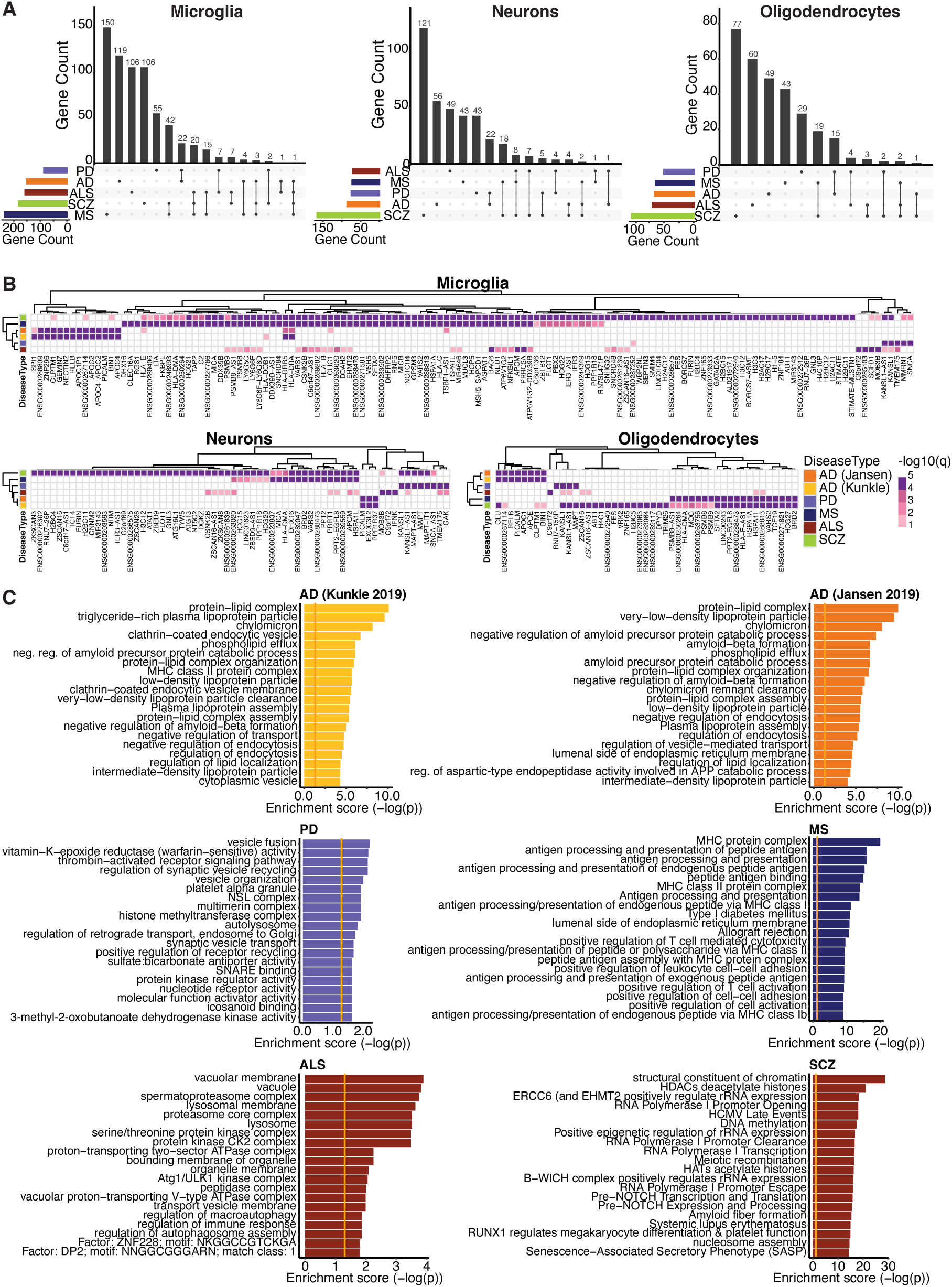
Microglia disease-risk genes impacted disease-specific pathways. **A)** UpSet visualization of unique and intersecting H-MAGMA disease-risk gene numbers between AD, PD (excluding 23andMe), MS, ALS and schizophrenia for each cell type. **B)** Heatmaps of H-MAGMA identified risk genes based on promoter-enhancer interactions from PLAC-seq data for AD, PD (excluding 23andMe), MS, ALS (p<5e-8) and schizophrenia (p<5e-12) for microglia, neurons and oligodendrocytes. Shown are H-MAGMA FDR corrected p-values (-log10(q)). **C)** Gene ontology pathway analysis of microglial risk genes identified by H-MAGMA for AD, PD (excluding 23andMe), MS, ALS, and schizophrenia; shown are top 20 pathways. SCZ, schizophrenia.

We next assessed specific cellular and biological pathways associated with microglia GWAS-risk genes for each disorder using gene ontology (GO) analysis. GO pathways linked to GWAS-risk genes were mostly unique for each neurodegenerative condition (**Fig. 3c**). This is consistent with the observation that most disease-risk genes were unique to each GWAS (**Fig. 3a, b**). The top GO pathways associated with microglial AD-risk genes included lipoproteins, amyloid processing and endocytosis (**Fig. 3c, Supplementary Table 2**) compared to neuronal and oligodendrocytes AD-risk genes which were associated with amyloid and tau protein catabolic processes (**Supplemental Fig. 3, 4**). PD microglial-risk genes were associated with the endolysosomal/autolysosomal pathways, synaptic vesicles and epigenetic signaling (**Fig. 3c, Supplementary Table 2**). Whereas substantia nigra gliosis, epigenetic signaling and synaptic vesicle pathways were evident in neuronal PD-risk genes, reinforcing the vulnerability of the midbrain in PD (**Supplemental Fig. 3**). Both microglia and oligodendrocyte MS risk-genes were associated with MHC protein complexes, autoimmunity, and antigen presentation and processing (**Fig. 3c, Supplemental Fig. 4, Supplementary Table 2**). Risk genes assigned to the MHC Class II complex were also associated with AD and PD, as well as MS (**Fig. 3b**). ALS exhibited associations with vacuoles and kinases, while also sharing pathways with PD related to lysosomes and autophagosomes (**Fig. 3c, Supplementary Table 2**). Microglial-associated GO pathways for schizophrenia GWAS-risk genes were distinct from the neurodegenerative disorders and primarily included epigenetic and gene regulatory pathways (**Fig. 3c, Supplementary Table 2**). Neuronal GWAS schizophrenia risk genes were primarily implicated in synaptic processes (**Supplemental Fig. 3**). Collectively, pathway analysis confirmed the observation from gene set overlaps, indicating that microglial risk genes and associated biological pathways are mostly disease-specific.

## Discussion

Incorporation of enhancer-to-promoter interactomes for microglia, neurons and oligodendrocytes with GWAS summary statistics enabled us to identify the cell types and genes associated with the genetic risk of brain disorders. Partitioned heritability analysis highlighted microglia as an important cell type underlying genetic susceptibility across multiple neurodegenerative conditions. Accordingly, enhancer-to-promoter interactomes identified the greatest number of predicted risk genes in microglia for AD, PD, MS and ALS. Previous studies have shown both the importance of active regulatory regions (63, 64) and that AD GWAS-risk is associated with gene regulatory regions in microglia (1, 41, 42, 56), as well as monocytes and macrophages (43, 65). MS is an autoimmune condition where the immune system attacks the myelin sheath surrounding neurons (66) and MS genetic risk genes have been associated with the peripheral immune system and microglia (67). ALS is a motor neuron disease that has been linked to aberrant inflammation (68), although GWAS risk for ALS has been primarily attributed to neuronal cell types (32). The genetic risk of PD using single-cell gene expression analysis has identified dopaminergic neurons and oligodendrocytes as cell types that express PD risk genes (69, 70). Interestingly, PD GWAS risk was found to be enriched in microglia and monocyte chromatin accessibility regions (71), although equivalent epigenetic datasets for dopaminergic neurons are lacking. In summary, chromatin interactions in microglia showed the strongest heritability enrichment and revealed the most risk genes across all neurodegenerative disorders. Despite this commonality, microglia genetic-susceptibility genes identified using H-MAGMA were associated with pathways that were disease-specific.

AD genetic risk in microglia was associated with lipoproteins, amyloid processing, endocytosis and MHC class II. The lipid-protein complex and lipoprotein pathways included the apolipoprotein genes *APOE*, *APOC1*, *APOC4-APOC2*, *APOC2* and *APOC4*, with APOE being the strongest common genetic determinate of sporadic AD (72). Amyloid processing pathways included the ABC transporter *ABCA7*, vesicle-associated genes such as *PICALM*, *BIN1* and *SORL1*, and protein cleavage genes such as *ADAM10* and *APH1B*. The endosome/endocytosis-associated AD risk genes *USP6NL*, *CNN2*, *RIN3*, *RAB8B* and membrane-associated genes such as *SPPL2A*, *STX4* may contribute to amyloid processing, although this remains to be fully explored. Rare loss of function variants for *ABCA7* and *SORL1* have also been implicated in increased AD risk (73, 74). The MHC class II complex was associated with AD risk and was mostly driven by the HLA locus (*HLA-DQB1*/*HLA-DRB1*/*HLA-DRB5*/*HLA-DRA/HLA-E*), as well as immune response genes such as *INPP5D*. Microglia mobility was implicated by AD risk genes such as the aggrecan protease *ADAMTS4* and cell adhesion molecule *CASS4*. Many AD risk genes were also implicated across pathways, for example, the low-density lipoprotein receptor, *SORL1*, recycles amyloid precursor protein out of endosomes (75).

PD risk genes in microglia were associated with endo-lysosomal pathways, as previously implicated in a non-cell-type-centric manner for PD (76). These included lysosomal-associated genes *LRRK2*, *RAB29* and *PLEKHM1*, as well as membrane fusion genes such as *STX4*, *TMEM175*, *VPS37A* and the familial PD gene *SNCA* (alpha-synuclein). The PD risk gene *ARHGAP27* has also been implicated in endocytosis (77). Histone modifications were associated with PD risk in microglia through histone lysine methylation (*SETD1A* and *FAM47E*) and acetylation (*KAT8* and *KANSL1*). Microglia and immune homeostasis, mobility and migration were linked to PD genetic risk through association with the purinergic nucleotide receptors *P2RY12* and *P2RY13*. Additional genes of interest are the vitamin K epoxide reductase *VKORC1*, the platelet-associated gene *MMRN1* and the kinases *DGKQ* and *CCNT2*. PD has been linked to mitochondrial dysfunction through familial mutations such as *PINK1* and *PARK7* (78) and environmental factors such as pesticides (79). The contribution of common PD-risk variants to mitochondrial function is less represented, however, we identified NADH:ubiquinone oxidoreductase complex assembly factor 2, *NDUFAF2*, the branched-chain keto acid dehydrogenase kinase, *BCKDK* and the G-protein-coupled receptor for succinate, *SUCNR1*, (citric acid cycle intermediate) as PD risk genes. BCKDK is localized to mitochondria and *BCKDK* mutations lead to dysregulated branched-chain amino acids and have been associated with Maple Syrup Urine Disease (MSUD) with links to Parkinsonism (80, 81). The microglia PD risk genes *LRRK2*, *SNCA,* and *TMEM175* have also been linked to rare coding mutations in PD patients (82, 83, 84).

MS-risk genes are mostly associated with T cell signaling and antigen presentation and processing, consistent with previous findings (85). A broader set of *HLA* genes were implicated in MS risk and genes linked to antigen presentation that were not identified in AD such as the ABC transporters *TAP1* and *TAP2*, TAP-binding protein *TAPBPL*, and the MHC Class I and Class II-associated genes (*MICB*, *CIITA*). Additional risk genes implicated in antigen processing were heat shock proteins (*HSPA1B*, *HSPA1A* and *HSPA1L*) and the ubiquitin ligase *MARCHF1*. MS-risk genes associated with immune activation included Tumor Necrosis Factor (*TNF*) and TNF receptor family members *TNFRSF1* and *CD27,* negative regulation of cytokines (*SOCS1* and *VSIR*), interleukin signaling (*IL12RB1*) as well as other immune signaling molecules such as *AIF1* (also known as IBA1), *BCL10* and *PTPRC*. Interestingly, several chromatin-related risk genes were identified including *CORO1A* and the lysine acetyltransferase *KAT8*.

Pathways for ALS risk genes were mostly associated with vacuole-related terms, as well as autophagy and the lysosome. These included vacuole-associated channels and transporters *ATXN3* (spinocerebellar ataxia-3), *CLCN3*, *SLC12A4*, *TMEM175* and lysosomal-associated proteins such as *TPP1*, *KICS2*, *NEU1*, *TM6SF1*, as well as the guanine nucleotide exchange factor *C9orf72*, iduronidase *IDUA*, formin binding protein *FNBP1* and the vacuolar ATPase *ATP6V1G2*. The proteasomal genes *PSMB8*, *PSMB9* and *PSMB10* were identified as MS-risk genes, with an isoform of *PSMB8* being linked to P-body formation in MS lesions (86). Several kinases were identified besides *C9orf72*, including *TBK1* and *CSNK2B*. Repeat expansions in C9orf72 and mutations in *TBK1* have established associations with both ALS and frontotemporal dementia (FTD) (87, 88).

The assignment of cell types to genetic risk and the identification of target genes depends on cell type epigenomic and chromatin interactome profiling. This has been performed for a limited number of cell types and chromatin conformation data has mostly been generated for non-dementia cases. Recent gene expression studies have implicated vascular cell types in the genetic risk for AD (89, 90, 91). Furthermore, the expression of AD risk has been reported to be differentially enriched in microglia substates (92). These examples highlight the need for epigenomic and chromatin conformation analysis of rare cell types and substates across disease conditions. However, our current analysis reinforces the genetic causative role of microglia in age-related brain conditions and offers biological insights into their involvement in various neurodegenerative disorders.

## Data and code availability

Code is available: https://github.com/aydanasg/cell_hmagma.

PLAC-seq, H3K27ac ChIP-seq, H3K4me3 ChIP-seq and ATAC-seq datasets were taken from (1) and processed data is available: https://github.com/nottalexi/brain-cell-type-peak-files.

## Author Contributions

AA contributed to investigation, methodology, project administration, formal analysis, data curation, visualization and writing-original draft. RMY contributed to data curation, supervision, writing and editing. SJM contributed to software, supervision, writing and editing AN contributed to conceptualization, methodology, resources, supervision and writing-original draft.

## Acknowledgements

We thank Dr Hyejun Won for the discussions on H-MAGMA. We thank the Skene, Marzi, Johnson and Nott groups for advice and helpful discussion, in particular, Alan Murphy, Dr Brian Schilder, Dr Kitty Murphy and Dr Alex Haglund.

## Materials and Methods

### PLAC-seq datasets

PLAC-seq data for human microglia, neurons and oligodendrocytes (1) was pre-processed by Nott et al., 2019 (1). PLAC-seq data was generated using epilepsy resections of the frontal, parietal and temporal cortex of seven individuals aged 5 months to 17 years. Chromatin interactions were 5 kb resolution and were anchored to promoters using chromatin immunoprecipitation of the histone modification H3K4me3 (1).

### Classification of PLAC-seq interactions

PLAC-seq chromatin interactions were classified as i) promoter-to-enhancer; ii) promoter-to-promoter; iii) promoter-to-ATAC; iv) promoter-to-promoter/enhancer; v) promoter-to-other; vi) H3K4me3-to-H3K4me3; vii) H3K4me3-to-other; and viii) other interactions for microglia, neurons and oligodendrocytes. ‘Promoter’ were classified as PLAC-seq bins that overlapped with H3K4me3 and H3K27ac regions within 2,000 bp of the nearest TSS and ‘enhancer’ were classified as PLAC-seq bins that overlapped H3K27ac regions distal to TSS as defined by Nott 2019 (1); promoter/enhancer were classified as PLAC-seq bins that overlapped both promoter and enhancer regions; ‘H3K4me3’ were PLAC-seq bins that overlapped H3K4me3 regions distal from TSS; ‘ATAC’ were PLAC-seq bins that overlapped chromatin accessibility regions that were devoid of H3K4me3 and H3K27ac; ‘other’ were PLAC-seq bins that did not overlap with H3K4me3, H3K27ac or chromatin accessibility regions (1). To identify the number of enhancers interacting with each promoter and number of promoters interacting with each enhancer, cell type PLAC-seq bins were overlapped with active promoter and active enhancer regions.

### GWAS datasets

The following GWAS summary statistics were used in this study were downloaded from EBI’s GWAS catalogue (https://www.ebi.ac.uk/gwas/) and were of European ancestry:

AD (Jansen 2019) (GCST007320): n= 71,880 cases and 383,378 controls (29);

AD (Kunkle 2019) (GCST007511): n = 21,982 cases and 41,944 controls, Stage 1 (28);

PD (Nalls 2019) (GCST009325): n = 33,674 cases and 449,056 controls (excluding 23andMe) (30);

MS (Andlauer 2016) (GCST003566): n = 4,888 cases and 10,395 controls (34);

ALS (van Rheenen 2021) (GCST90027164): n = 27,205 cases and 110,881 controls (32);

schizophrenia (Trubetskoy 2022) (GCST90128471): n = 53,386 cases and 77,258 controls (33).

### Quality control of GWAS summary statistics

GWAS summary statistics were standardised and underwent quality control steps before running H-MAGMA. GWAS summary statistics were filtered using format_sumstats function in “MungeSumstats” package (version 1.6.0, available on Bioconductor) in R (version 4.2.1) (93). Summary statistics had the following imputation quality: AD (Jansen 2019) >0.91; AD (Kunkle 2019) >0.4; PD > 0.8; MS ≥0.8; ALS >0.95; schizophrenia (INFO>0.9).

### H-MAGMA

Annotating genetic variants to target genes was performed using H-MAGMA (52, 94). H-MAGMA input files provide the background profile of gene-SNP associations based on chromatin interaction data. To generate cell type-specific promoter-enhancer profiles, 1) chromatin interaction data from PLAC-seq for microglia, neurons and oligodendrocytes, and 2) reference data for SNPs (22665064 million SNPs) from Phase 3 of 1,000 Genomes for European ancestry were used (genome Build 37) (https://ctg.cncr.nl/software/magma). Exonic and promoter SNPs were directly assigned to target genes based on genomic location using a gene model Gencode v41 (https://www.gencodegenes.org/human/release_41lift37.html) (95). Promoters were defined as 1.5kb upstream and 500bp downstream of the TSS of each gene isoform. Intronic and intergenic SNPs were assigned to cognate genes based on cell-type chromatin interactions (see PLAC-seq datasets) with promoters and exons (52). Intronic and intergenic SNPs were filtered to enhancer SNPs by overlapping with cell-type enhancer regions (1). To investigate disease enrichment in active chromatin interactions, significant cell-type specific chromatin interactions with FDR-corrected p-value cut-off of 0.01 were filtered to interactions with promoters in at least one end by overlapping cell-type promoter regions (1). Filtered chromatin interactions were overlapped with Gencode 41 exon and promoter coordinates to identify exon-based and promoter-based interactions (52, 94). To determine whether enhancer or promoter/exon SNPs were driving the disease enrichment of genes, H-MAGMA input files were generated either with promoter/exon SNPs or enhancer SNPs only. H-MAGMA outputted genes with an FDR-corrected p-value <0.05 were selected for downstream analysis.

### MAGMA

MAGMA analysis pipeline was used to run the H-MAGMA cell type-specific gene level association with a disease (53). The association was established using the default “SNP-wise mean” gene analysis model, which is a test of mean SNP association using the sum of squared SNP Z-statistics as a test statistic. In brief, SNP-level p-values from GWAS summary statistics were aggregated into gene-level p-values and a reference data set (1,000 Genomes European panel) was used to account for linkage disequilibrium between SNPs. Since some of the GWAS summary statistics used in the study are SNP meta-analysis results, individual sample sizes per SNP may have significant variation and may affect the gene test-statistic results. Therefore, if available, individual sample sizes per SNP were used (ncol modifier in –pval parameter in MAGMA). The analysis was run as follows: magma --bfile g1000_eur --pval <GWAS summary statistics> use=SNP,P ncol=NSUM --gene-annot <INPUT annotation file> --debug set-spar=tmp_snps_used --out <OUTPUT file>.

### Partitioned heritability (sLDSC regression)

Partitioned heritability using sLDSC regression analysis was used to identify brain cell type annotations that were enriched for heritability of AD, PD (excluding 23andMe), MS, ALS and schizophrenia (LDSC version 1.0.1) by functional category while controlling for 97 annotation categories of the full baseline model (model version 2.2) (96). Cell type annotations per functional category were run jointly. Functional categories included cell type 1) total PLAC-seq bins, 2) promoter and enhancer PLAC-seq bins, 3) promoters PLAC-seq bins, and 4) enhancer PLAC-seq for microglia, neurons and oligodendrocytes. Baseline model LD scores, standard regression weights, and allele frequencies that were used were built from 1000 Genomes Phase 3 for European population. The enrichment P-values were FDR multiple testing corrected for the number of GWAS studies and number of cell types using Benjamini-Hochberg correction method. Disease enrichment was considered insignificant if the coefficient z-score was negative. Cell type annotations for all the functional categories were created using plink format .bed/.bim/.fam filesets of 1000 Genomes Phase 3 for European population and LD scores were computed based on a 1 centiMorgan (cM) window. Since the annotations were built on top of the baseline model, 1000 Genomes Phase 3 was used together with the HapMap3 SNPs. A quality control step of GWAS summary statistics was performed before LDSC analysis using munge_sumstats.py where SNPs had INFO <= 0.9, MAF <= 0.01 and N < 32290, were out-of-bounds p-values, strand-ambiguous, with duplicated IDs and alleles did not match Hap-Map SNPs. To prevent bias from variable imputation quality both between and within each GWAS study, all the GWAS SNPs were filtered to HapMap3 SNPs, as these SNPs are well imputed in most studies.

### EWCE

Expression weighted cell type enrichment (EWCE) analysis (v1.6.0) was used to identify cell type-specificity of the H-MAGMA outputted risk genes for each disease type (97). Single-cell RNA-seq data from mouse cortex and hypothalamus from Zeisel et al. (2015) study (58) was used to generate probability distribution associated with cell type-specific H-MAGMA outputted risk genes having an average level of expression within a cell type. Significant cell type-specificity was determined based on the p-value <0.05.

### GO analysis

Gene set enrichment analysis was performed on the list of H-MAGMA outputted significant risk genes identified per cell type to identify biological pathways at risk in each cell type for each disease. The R package “gprofiler2” (v0.2.1) was used for gene set enrichment, which contains data sources including Gene Ontology (GO), KEGG, Reactome, WikiPathways, miRTarBase, TRANSFAC, Human Protein Atlas, protein complexes from CORUM and Human Phenotype Ontology (98). Risk genes inputted into the analysis were filtered based on the FDR adjusted p-value<0.05 and were ordered based on the Z-score generated by the H-MAGMA. Identified pathways were also FDR corrected using p-value <0.05. For visualization, if pathways contained the same set of genes, the one with the highest FDR corrected p-value was included in the bar plots.

---

**Supplementary Figure 1.**
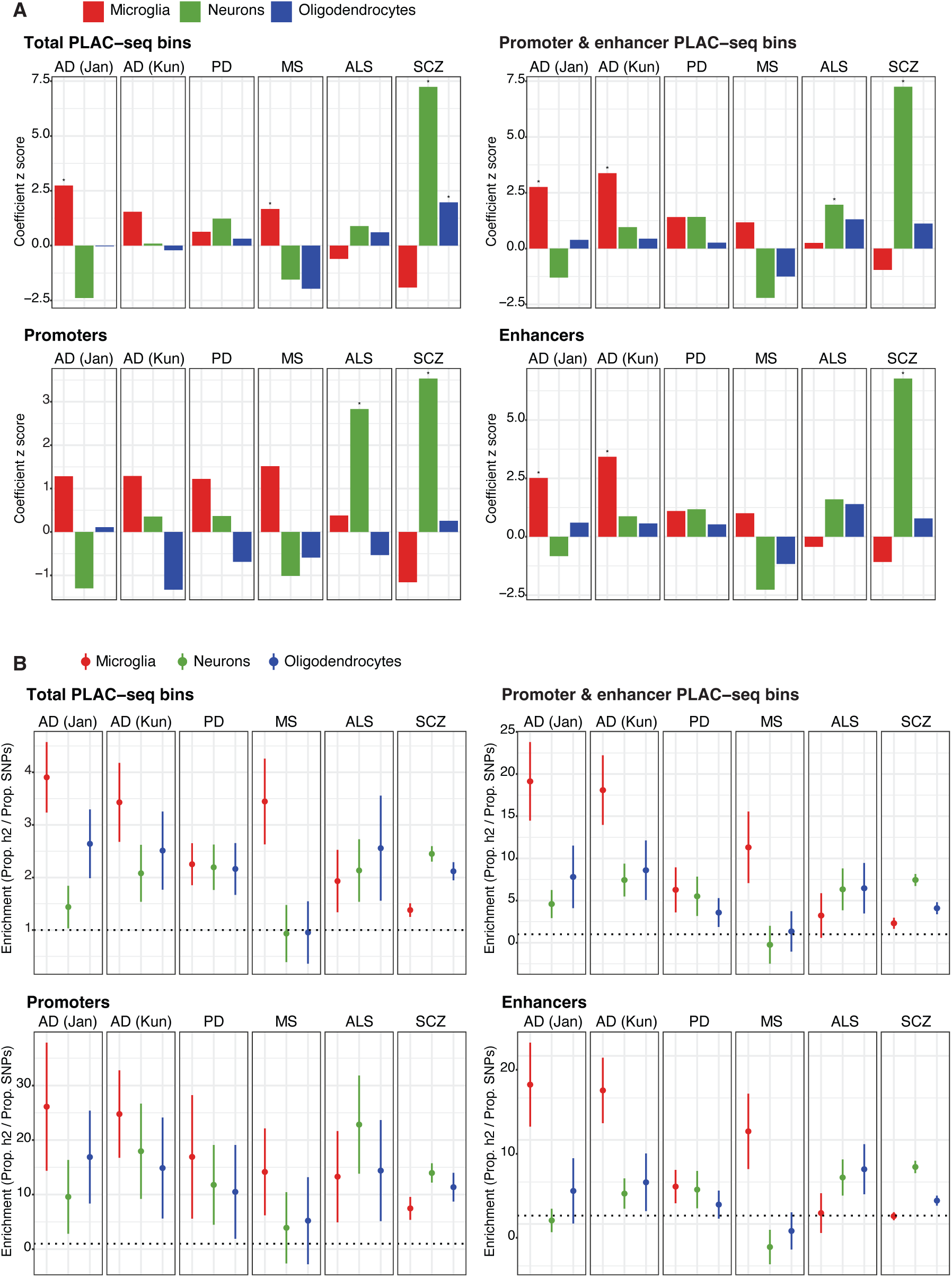
LDSC coefficient z-scores and enrichment values. **A)** Partitioned heritability sLDSC coefficient z-scores for i) total PLAC-seq bins (ii) promoter and enhancer PLAC-seq bins; iii) all promoters and iv) all enhancers for microglia, neurons and oligodendrocytes in AD, PD (excluding 23andMe), MS, ALS, and schizophrenia. *transformed coefficient p-values < 0.05. **B)** Partitioned heritability sLDSC enrichment values defined as the ratio of the proportion of heritability to the number of SNPs (Prop. h2 / Prop. SNPs) for i) total PLAC-seq bins (ii) promoter and enhancer PLAC-seq bins; iii) all promoters and iv) all enhancers for microglia, neurons and oligodendrocytes in AD, PD (excluding 23andMe), MS, ALS, and schizophrenia. The grey dotted line represents the cutoff for enrichment (1). Error bars represent standard error. SCZ, schizophrenia.

**Supplementary Figure 2.**
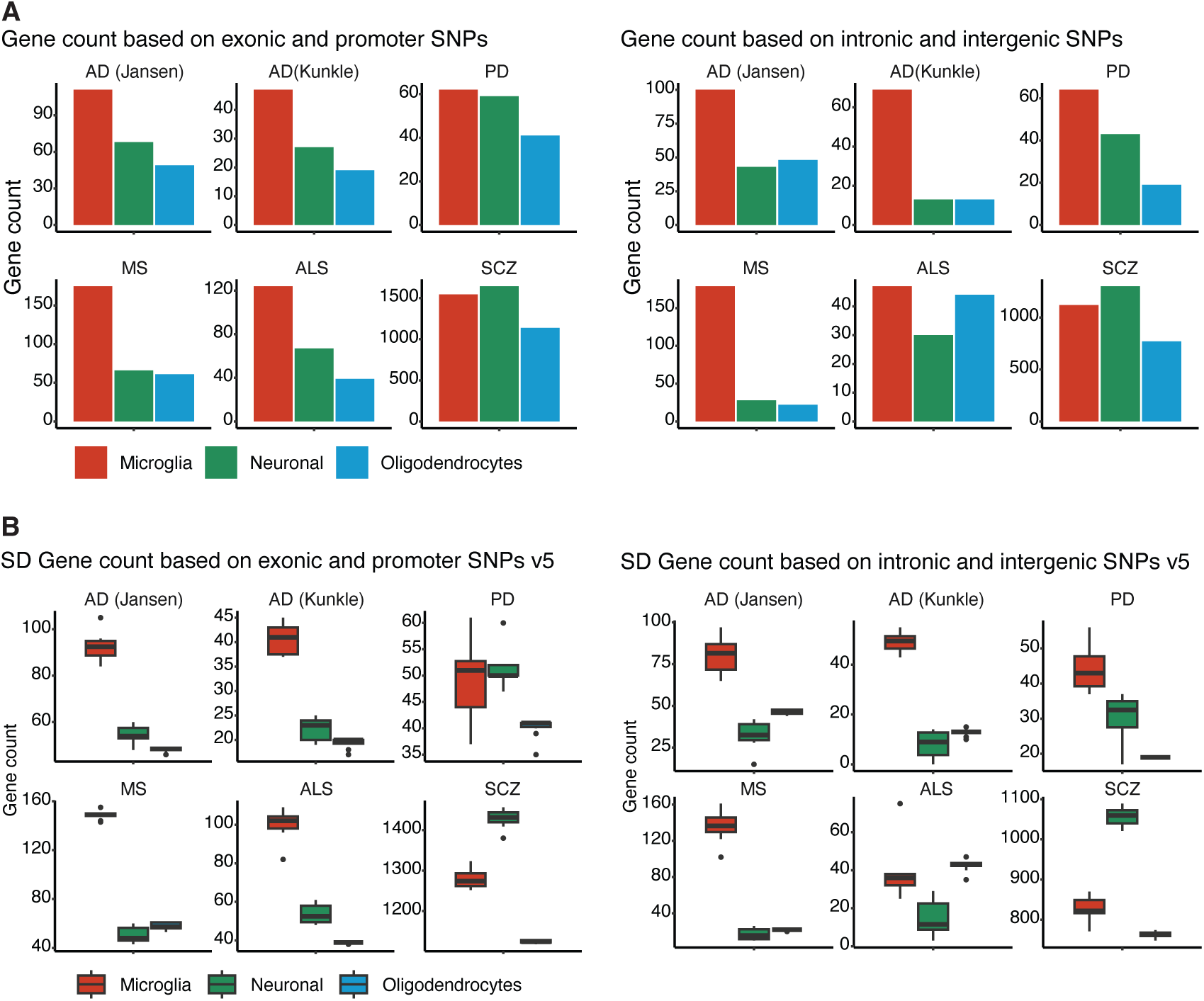
H-MAGMA disease risk genes identified using PLAC-seq interactions overlapping SNPs subset to either genes or enhancers only. A) The number of disease-risk genes identified in microglia, neurons and oligodendrocytes using H-MAGMA and GWAS for AD, PD (excluding 23andMe), MS, ALS, and schizophrenia using SNPs overlapping PLAC-seq bins at i) exon and promoters only (left) or at ii) enhancer regions only (right). B) Chromatin interactions were randomly sampled down 10 times to 60,000 interactions and the number of disease-risk genes were identified in microglia, neurons and oligodendrocytes using H-MAGMA and GWAS for AD, PD (excluding 23andMe), MS, ALS, and schizophrenia using SNPs overlapping PLAC-seq bins at i) exons and promoters only (left) or at ii) enhancers only (right). Dunn’s test (non-parametric) between cell types within each group: i) exons and promoters only: AD (Jansen 2019): microglia-oligodendrocytes (**), PD: microglia-oligodendrocytes (**), neurons-oligodendrocytes (***); MS: microglia-neurons (****), microglia-oligodendrocytes (**); ALS: microglia-neurons (*), microglia-oligo (****), neurons-oligo (*); schizophrenia: microglia-neurons (****), microglia-oligo (*****) and ii) enhancers only: AD (Jansen 2019): microglia-neurons (****), microglia-oligo (**); PD: microglia-neurons (**), microglia-oligo (****), neurons-oligo (*); MS: microglia-neurons (****), microglia-oligo (**); ALS: microglia-neurons (**), neurons-oligo (****); schizophrenia: microglia-neurons (*), microglia-oligo (*), neurons-oligo (****). *p <0.05, **p<0.01, ***p<1e-4, ****p<1e-6. SCZ, schizophrenia.

**Supplementary Figure 3.**
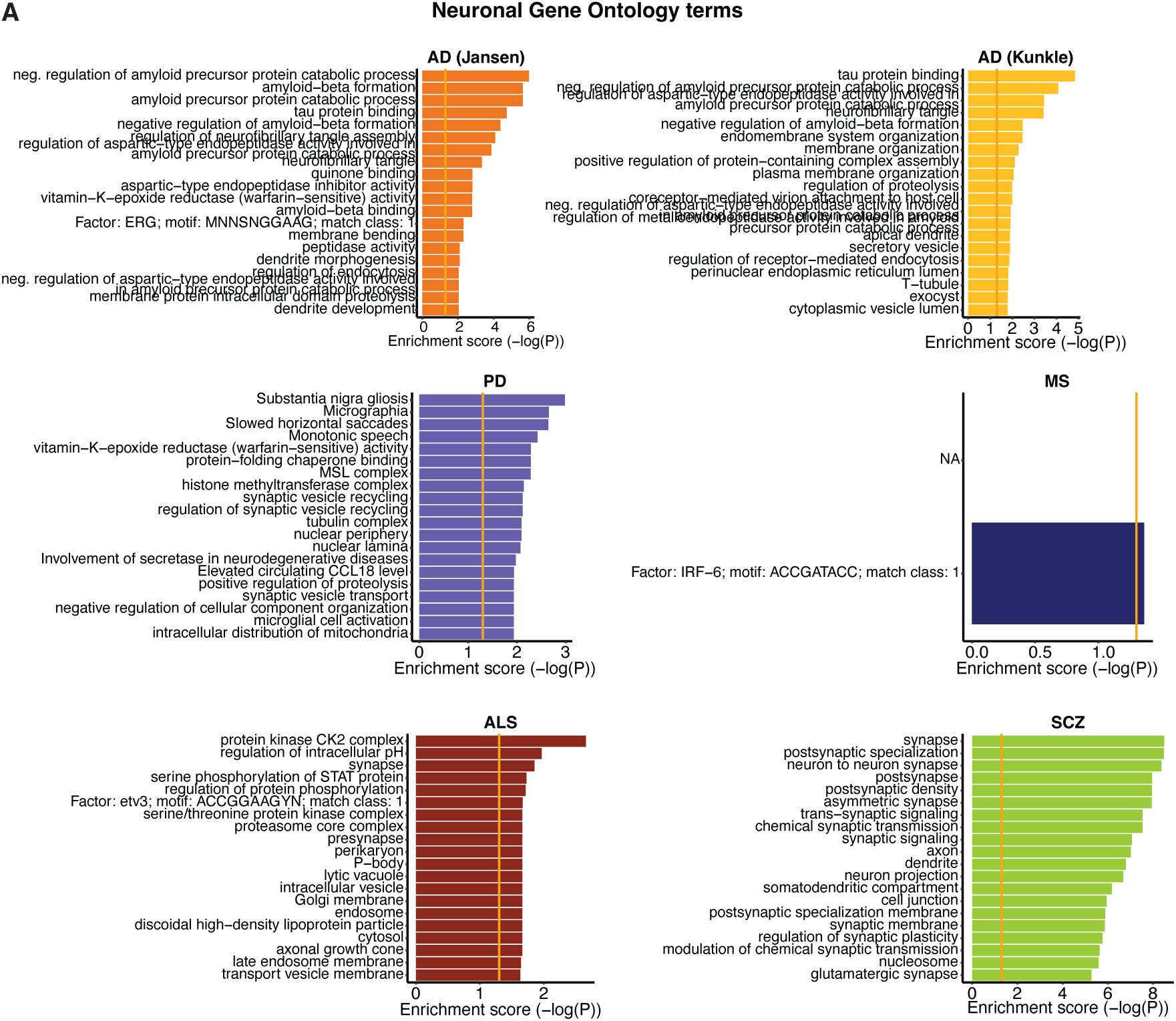
Gene ontology pathways for neurons across diseases. Gene ontology pathway analysis of neuronal risk genes identified by H-MAGMA for AD, PD (excluding 23andMe), MS, ALS, and schizophrenia; shown are the top 20 pathways. SCZ, schizophrenia.

**Supplementary Figure 4.**
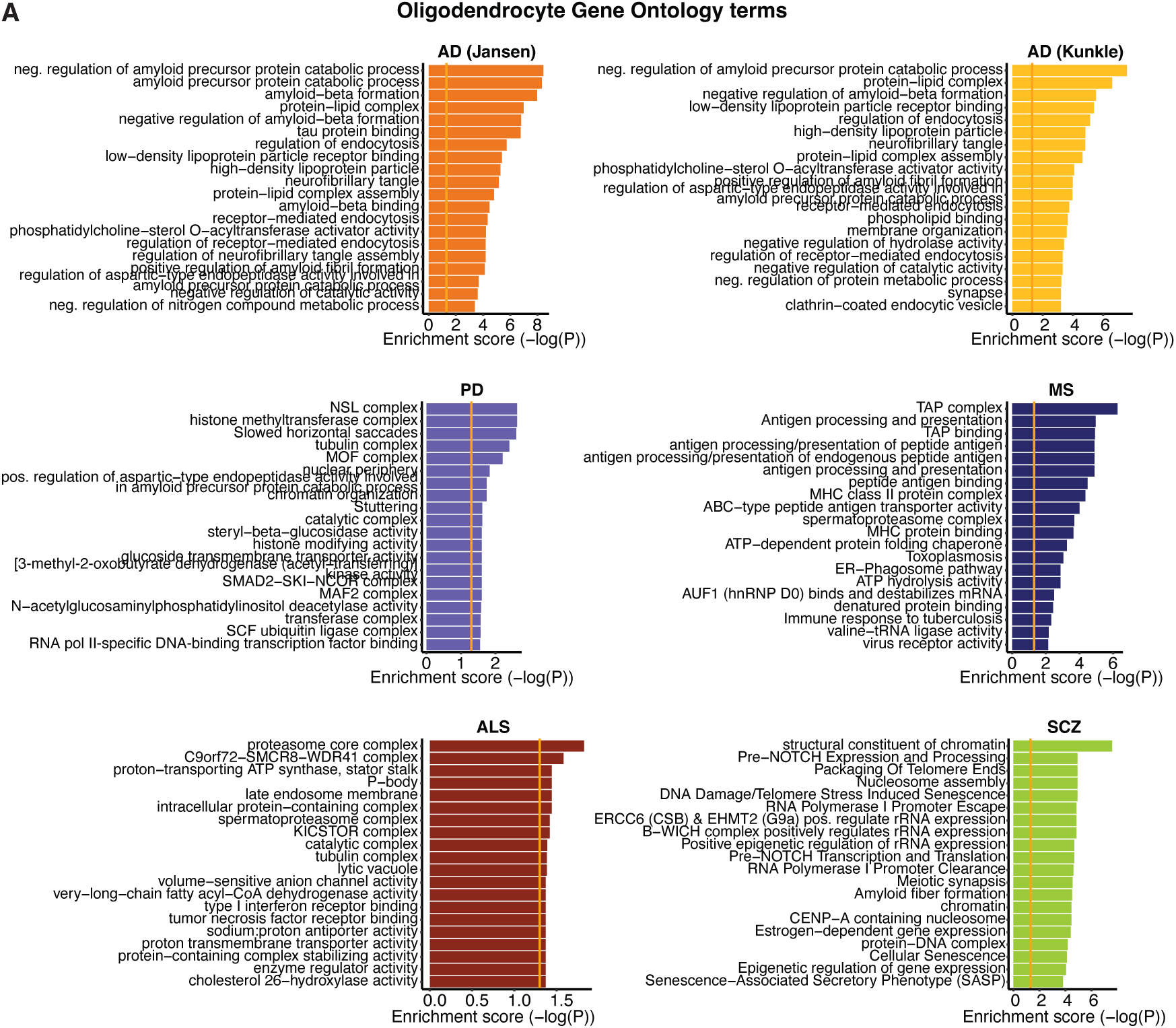
Gene ontology pathways for oligodendrocytes across diseases. Gene ontology pathway analysis of oligodendrocyte risk genes identified by H-MAGMA for AD, PD (excluding 23andMe), MS, ALS, and schizophrenia; shown are the top 20 pathways. SCZ, schizophrenia.

